# Impact of bilayer composition on the dimerization properties of the Slg1 stress sensor TMD from a multiscale analysis ^†^

**DOI:** 10.1101/2022.07.27.501806

**Authors:** Fabian Keller, Azadeh Alavizargar, Roland Wedlich-Söldner, Andreas Heuer

**Affiliations:** Institut für Physikalische Chemie, Corrensstraße 28, Münster, Germany; Institut für Zelldynamik und Bildgebung, Von-Esmarch-Straße 56, Münster, Germany

## Abstract

Mutual interactions between the transmembrane domains of membrane proteins and lipids on the bilayer properties has gained major interest. Most simulation studies of membranes rely on the Martini force field, which has proven extremely helpful in providing molecular insights into realistic systems. Accordingly, an evaluation of the accuracy of Martini is crucial to be able to correctly interpret the reported data. In this study, we combine atomistic and coarse-grained Martini simulations to investigate the properties of transmembrane domains (TMDs) in a model yeast membrane. The results show that the TMD binding state (monomeric, dimeric with positive or negative crossing angle) and the membrane composition significantly influence the properties around the TMDs and change TMD-TMD and TMD-lipid affinities. Furthermore, ergosterol (ERG) exhibits strong affinity to TMD dimers. Importantly, the right-handed TMD dimer configuration is stabilized via TMD-TMD contacts by addition of asymmetric anionic PS. The CG simulations corroborate many of these findings, with two notable exceptions: a systematic overestimation of TMD-ERG interaction and lack of stabilization of the right-handed TMD dimers with the addition of PS. Atomistic simulation results suggest that a meaningful comparison of dimer formation and experimentally-determined network factor may require to additionally take into account the precise conformation and thermodynamic relevance of multimeric TMD clusters.

## 1 Introduction

The proper functioning of membrane proteins is critical for a cell to communicate with its environment. Therefore, understanding how membrane proteins, the active constituents in cell membranes, take effect and how their activity can be modified is crucial for, e.g., rational drug design.^1–5^ While the transmembrane domains (TMDs), especially that of bitopic proteins, were considered to be the mere anchors in the cell membranes, the relevance of TMDs in several processes has become evident in the recent decade: TMDs were found to be linked to intermembrane signal transduction, vesicle fusion, or a means of particle transport through the membrane. The predominant structural motif for TMDs is a *α*-helix.^6–8^ Their sequences are conserved, in particular at one side.^9^ In fact, most TMD sequences are highly adapted to the properties of their target membranes.^7^

The functionality of membrane proteins often depends on their correct position within cell membranes^8,10–13^ and this position is controlled via the interaction with the local lipid environments.^10,12,14,15^ This interaction can occur via two modes. First, annular lipids form an annulus around the TMD that differs from the bulk properties in composition, their chain order and dynamics. ^11,13^ Their binding strength in this mode depends on the TMD length (hydrophobic mismatch) and thickness, which determines the TMD’s affinity for ordered versus disordered lipids.^11,16^ The adaptation of the cell to alter the TMD’s affinity to particular lipids is the basis for organellar targeting and membrane domain formation. ^7^ Second, non-annular or structural lipids can interact with specific binding motifs on TMDs. For cholesterol (and likely also for the corresponding sterol in Yeast cells, ERG), the CRAC, CARC, and CCM binding motifs were identified as specific interaction sites^17–19^. These interactions between cholesterol and the binding motifs were shown to be transient with a rapid molecule exchange.^18,20,21^ Besides these motifs, positively charged amino acids (e.g., Arg, Lys) in the juxtamembrane position were found to interact strongly with negatively charged lipids, concentrated on the cytosolic leaflets of biological membranes and may in turn control the surrounding lipid composition. ^22–24^

Besides the correct localization of membrane proteins in the cell membranes, the interaction of the membrane proteins, especially their TMDs, determines their activation state, as most bitopic TMDs are activated via dimerization.^25,26^ The GxxxG motif is associated with strong TMD-TMD binding, where x stands for hydrophobic residues with short side-chains.^27,28^ In the canonical motif, glycine residues form a groove with a flat contact surface. Small residues (Ala, Thr, Cys, Ser) may replace glycine, and thus, the motif can be generalized to SmxxxSm (Sm for small residue). On the other hand, lipid contacts may influence the TMD binding.^29^ Multiple studies have reported that the lipid environment influences the strength of the TMD-TMD binding^2,26^ and especially cholesterol was shown to modify the binding strength of TMD dimers. ^1,3,4^

Of general interest is the association process of TMDs and in particular, as the elementary step, the formation of TMD dimers and their structural properties. Since the mechanism of TMD association is a slow process, which has been practiced using atomistic (AA) simulations together with enhanced sampling techniques^30–34^, coarse-grained (CG) Martini MD simulations are an appropriate alternative to achieve relevant time scales. Subsequently, the CG configurations can then be mapped on an AA representation to achieve a more microscopic perspective. Examples in the context of dimerization of TMDs can be found in refs. ^33,35–37^.

Recently, we have studied in a joined project, involving experiments and simulations, the yeast cell wall stress sensor Slg1 as a representative membrane protein TMD^38^. The sequence and structure of a dimer of the Slg1-TMD are depicted in figure 1. Slg1 is a group I protein (N-terminal facing to cell exterior) and features two specific interaction sites: GxxxG-like motifs can be found in the extracellular leaflet and in the cytosolic leaflet, the charged residues, histidine and arginine, lie in the head group region of the bilayer. The simulations were conducted using the CG Martini 2 force field and the bilayers were chosen with symmetric and asymmetric (additionally containing charged PS lipids) leaflet compositions. The lipid compositions were chosen to model the yeast PM (corresponding to the work of Ejsing and coworkers^39^). Of particular interest was the change of the free energies of dimerization upon addition of the PS lipids. In agreement with the network factor in the corresponding microscopy experiment, also reflecting the association behavior of TMDs, we observed no appreciable effect of the addition of PS lipids. Furthermore, we observed a strong interaction of ERG with the GxxxG motifs.

**Fig. 1.**
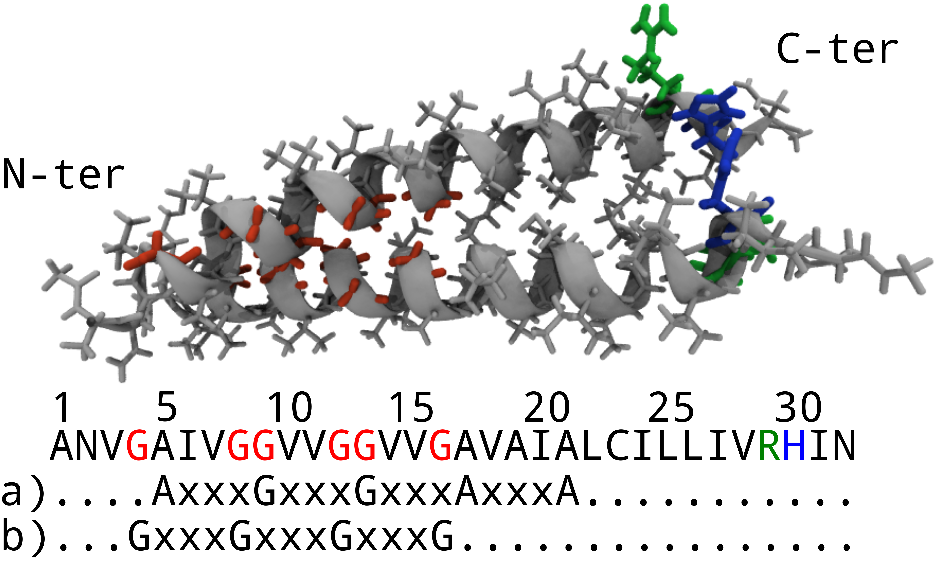
Structure of the Slg1 TMD dimer (top) used in this simulation study. Glycine, arginine, and histidine residues are highlighted in red, green, and blue, respectively. The one-letter coded protein sequence (bottom) of the used TMD is shown with the glycine-zipper(-like) motifs a) and b).

In this work, we present AA simulations of Slg1 dimers and monomers, obtained from backmapping of the corresponding CG configurations. Our aim is threefold. Firstly, we characterize the structural properties of TMD dimers and TMD monomers on the AA level, thereby exploring the molecular fidelity on the atomistic scale. Of particular interest is their respective impact on the surrounding lipids and, reversely, the impact of the lipids on the dimer properties. This involves, in particular, the specific role of the GxxxG-motif as well as the binding of ERG and PS with the TMD dimer, reflecting their impact on the dimer properties. Secondly, we analyse whether the equilibrated CG dimer configurations and AA dimer configurations are sufficiently close to each other. For this purpose we study the time evolution of backmapped (from CG to AA) dimer configurations. To ensure sufficient sampling of the dimer configurations in atomistic resolution, we simulated the systems containing dimers for over 6.5 *μ*s. Thirdly, we provide a careful comparison of the structural properties of the AA and CG configurations to highlight consistent but also non-consistent structural and dynamical features. It is known that Martini 2 tends to have too strong protein-protein interactions, which is supposed to be remedied by the newly published Martini 3 force field^40^. However, so far Martini 3 does not include a consistent parametrization of sterol molecules so that for the time being simulations of biologically relevant lipid mixtures still rely on the use of Martini 2. In total, new insight about the structural properties of TMD dimers is provided, highlighting the similarities and differences from simulations on different scales and the implications for the interpretation of the observed association behavior in microscopy experiments.

## 2 Computational details

### 2.1 Coarse-grained simulations

The CG simulations of the Slg1 TMD in membrane bilayers was part of our previous work^38^. Here we summarize the important aspects of the modeling procedure. The initial atomistic structure of this TMD was constructed using the SWISS-MODEL homology modeling web server.^41–43^ The two initial monomer structures were aligned in the z-direction at a distance of 4.5 Å from each other. Subsequently, the monomers were inserted in membranes with different compositions and solvated with water molecules, with the addition of ions to neutralize the system, using Martini maker as implemented in web-based CHARMM-GUI membrane builder. ^44^ The final systems contain two TMD domains and around 400 lipid molecules. Two systems were generated, which have the most similar lipid composition as the implemented experiments, as discussed in the previous work^38^. Both systems contain 20% ERG, one system is symmetric and additionally contains PC lipids, and the other system is asymmetric and it includes also PS lipids with the amount of 27% in the lower leaflet. For each lipid composition, two samples with a right-handed and left-handed dimer configuration were selected for further analysis. CG molecular dynamics (MD) simulations were performed using the Martini 2.2 force field to describe the interactions between the peptides, lipids and solvent molecules. ^45–47^ In Martini representation, on average 4 atoms along the associated hydrogens are represented by one interaction site (bead). Since the parameters for the sterol molecules are not yet available for the latest version of Martini and the membrane of Yeast contains a considerable amount of ERG, we used Martini 2.2 for our simulations.

All CG MD simulations were performed using Gromacs version 2019.^48,49^ To maintain the temperature at 310 K, the system was coupled to a Berendsen thermostat^50^ for equilibration and a velocity rescaling thermostat^51^ for production simulations. To control the pressure at 1 bar, a Berendsen pressure barostat for equilibration and a Parrinello–Rahman barostat^52^ were used with coupling constant of 12 ps and compressibility of 3 × 10^-4^ bar^-1^) for production simulations, using semi-isotropic coupling scheme. The non-bonded interactions were treated using a switch function from 0.0 to 1.1 nm for the electrostatic and from 0.9 to 1.1 nm for the Lennard Jones interactions. For equilibration, the input files provided by the CHARMM-GUI web server were used. Restraints were applied on the proteins and lipids and gradually reduced to zero. The simulation time for each system is 10 μs (corresponding to effective time of 40 μs), performed in the NPT ensemble using a time step of 20 fs. A detailed list of the compositions and simulations lengths of the CG simulations is specified in table S1^†^.

### 2.2 Atomistic simulations

For the atomistic simulations, three different TMD configurations were prepared and integrated into an asymmetric and a symmetric bilayer. The TMD configurations included a TMD monomer, a TMD dimer with negative crossing angle ***χ***(−), a TMD dimer with positive crossing angle ***χ*** (+). The bilayer structures with a TMD monomer were obtained from the CHARMM-GUI website ^53^ and the initial bilayer structures with TMD dimers were obtained from the CG simulations and transformed to atomistic resolution by using the Martini backmapping script (see section 2.3 for further discussions).

The TMDs were acetylated at the N-terminus and methylamidated at the C-terminus to avoid artifacts from unphysiological charges. The asymmetric bilayers comprised PS, PC, and ERG lipids in the cytosolic leaflet with a PL ratio of 1:6 (PS:PC) and 20% ERG. The extracellular leaflets contained only PC and ERG lipids. Each PL head group type included three different chain types with a ratio of (1:1:1), namely PO(18:1-16:0), YO(18:1-16:1), or DY(16:1-16:1).

The atomistic simulations were conducted using the CHARMM36 force-field^54^,^55^ and Gromacs 2018 ^56^,^57^. The systems were equilibrated using the 6-step established CHARMM-GUI parameter set gradually decreasing restraints on lipid head group positions and chain dihedral angles (see table S3^†^). The simulations of the dimers were run for at least 6 μs and simulations of the TMD monomers were at least run for 2μs. A detailed list of the compositions and simulations lengths of the atomistic simulations is specified in table S2^†^.

The TIP3P model was used as water model.^58^ The Nosé-Hoover algorithm was used to maintain the temperature at 300 K with a coupling constant of 1 ps, coupling bilayer and solvent separately. ^59^ To maintain the pressure at 1 bar the Parrinello-Rahman barostat was used with a semiisotropric coupling scheme, a coupling constant of 5 ps and compressibility of 4.5×10^-5^ bar^-1^.^52^ Particle mesh Ewald electrostatics were used with a real-space cutoff of 1.2 nm.^60^ The Lennard-Jones potential was shifted to zero between 1.0 and 1.2 nm, with a cutoff of 1.2 nm and the nonbonded interaction neighbor list was updated every 20 steps with a cutoff of 1.2 nm.

### 2.3 Backmapping of Martini Coarse-Grained bilayer structures

For the backmapping from CG to AA, representative structures within the CG simulations had to be selected. In the course of these simulations, the two monomers formed dimers after around 900 ns to 2 μs. For each system, two structures were selected in which right-handed and left-handed dimers have been formed with relatively low crossing angle values (24° and 22° for righthanded and 19° and 29° for left-handed, respectively, for the symmetric and asymmetric systems). We used the backward algorithm for the backmapping of the selected structures. ^61,62^ In this approach, atoms are placed alongside bonds between two CG beads. A subsequent geometric correction was made based on geometric rules to obtain correct double bond isomery and chiral centers. In some cases the backmapping process failed to produce correct stereochemistry for chiral centers of ERG molecules and PL double bonds. We, therefore, wrote a script to identify erroneous stereochemistry. Chiral centers were changed from right-to left-handed by switching planes of the hydrogen from one side of the plane spanned by the three other substituents to the other. Double bonds were switched from <u>trans</u> to <u>cis</u> by mirroring the positions of the =C**H** and the =C**CH**_2_. Subsequent energy minimizations provided robust results with correct stereochemistry.

## 3 Results

### 3.1 Characterization of the TMD-lipid environment

The sequence of the Slg1 TMD includes two important interaction sites: The GxxxG motifs, associated with strong TMD dimerization and ERG affinity, and the charged residues R/H associated with strong binding to charged lipids. We, therefore, calculated the enrichment of ERG, PC, and PS lipids around the TMDs. The enrichment describes the sum of contacts between a lipid type and all residues of a predefined group normalized by the total number of residue group-lipid contacts and the bulk concentration (relative fraction) of the lipid types in the corresponding leaflet. The resulting ratio is subtracted by 1. Hence, 0 indicates no enrichment, values >0 indicate enrichment, and values <0 indicate depletion. The data are averages over the two TMDs in the dimers and periods of 300 ns. In this definition residue-lipid contacts are formed if they had at least a pair of atoms within 0.4 nm (two times the typical water hydrogen bond length). In figure 2 we display the corresponding data around the TMDs in the cytosolic side in the asymmetric membranes (the corresponding data for the symmetric membranes is displayed in figure S3^†^), and in figure 3 we show the enrichment around the two motiffaces and the other faces on the extracellular side.

**Fig. 2.**
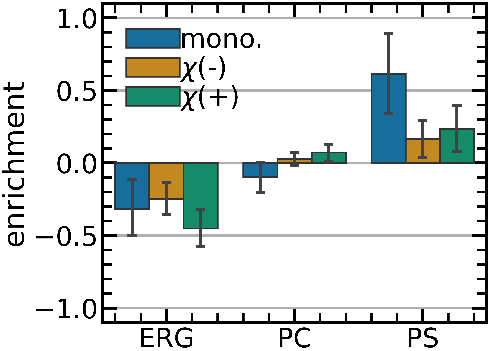
Enrichment of ERG, PC and PS lipids in the cytosolic leaflet of the different asymmetric bilayer setups (monomeric, or dimeric in righthanded ***χ***(−) and left-handed ***χ***(+) configuration) around the TMD. The bars indicate the 68% confidence interval (1 standard error).

**Fig. 3.**
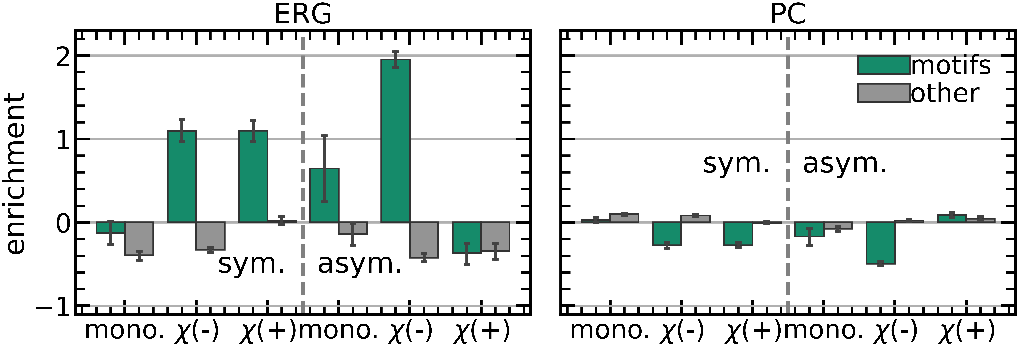
Enrichment of ERG and PC lipids in the extracellular leaflet of the different bilayer setups (symmetric vs. asymmetric, monomeric or dimeric in right-handed ***χ***(−) and left-handed ***χ***(+) configuration) around the motif faces a) and b) (green, see figure 1) and the other faces (gray). The enrichment is a measure for the agglomeration of a lipid type with respect to its bulk concentration (see the main text for the definition). The bars indicate the 68% confidence interval (1 standard error).

In the cytosolic leaflet, the TMD enrichment of PS provides insights into the effects of the charged residue R/H. Indeed, we see enrichment of PS near the TMD (figure 2). The PS enrichment is smaller than the enrichment of ERG at the motif faces (enrichments of >1 versus <0.5) and may even be negligible in the case of the dimers. PC lipids show no enrichment, and ERG shows depletion (similar to the “other” face in the extracellular leaflet). In the symmetric bilayers, the enrichment data indicates similar behavior (figure S3^†^).

The enrichment values of PC and ERG indicate distinct affinities to the TMD faces (figure 3). ERG has particularly enriched near the TMD motif faces of the dimers, except for the ***χ*** (+) dimer in the asymmetric bilayer. The lipid compositions appear to have a significant impact on the ERG affinities. In the asymmetric bilayers, the dimer configurations exhibit different enrichment profiles with profound enrichment at ***χ*** (−) and depletion at ***χ*** (+). However, in the symmetric bilayers, the enrichment is similar for both dimer configurations. Near the TMD monomers, the motif faces show no enrichment (symmetric bilayer) or some enrichment albeit with high uncertainty (asymmetric bilayer). PC lipids, on the other hand, show no enrichment whatsoever.

Indeed, ERG shows a high affinity to the GxxxG-motifs, consistent with the interaction of cholesterol with other TMDs^18,29,63–65^, especially around the dimers (except the ***χ***(+) dimer in the asymmetric bilayer), while the enrichment around the monomers is weak (asymmetric bilayer) or absent (symmetric bilayer). This likely arises from a similar effect as lipophobic effect, which is due to the unfavorable state of the lipids around the monomer as a result of the exposure of polar residues to the lipid phase^66,67^.

We characterized the TMD-ERG and TMD-PS bond strength by measuring the time a lipid neighbor remains attached to the respective TMD. The average residence times ***τ*** and the ratio of the averages and the corresponding standard deviations are depicted in figure 4. Only residence times longer than 5 ns and lipid contacts that were broken before the end of the simulations were considered. The fraction of the mean and standard deviation of the residence time enables us to distinguish between a homogeneous time distribution where the residence times follow simple exponential decay (***σ***(***τ***)/ < ***τ*** >= 1) or more complex behavior with some lipids having long residence times (s(***τ***)/ < ***τ*** >> 1).

**Fig. 4.**
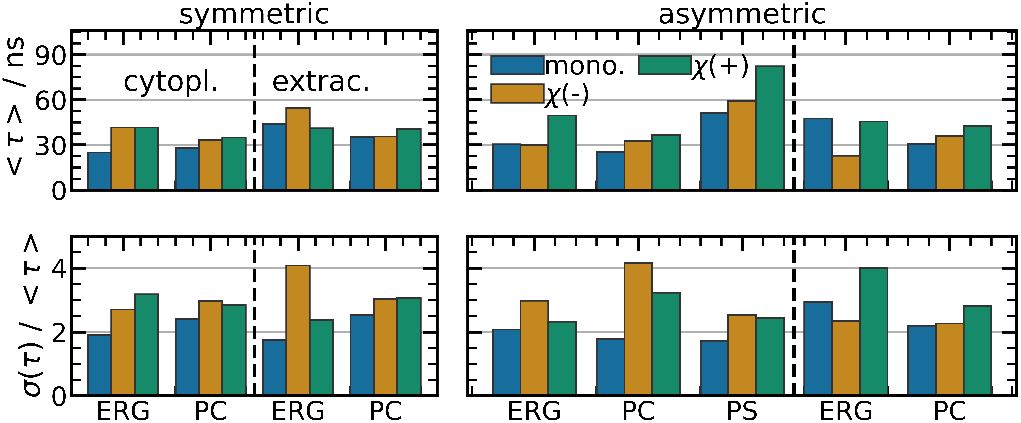
The average (top) residence times ***τ*** of lipid neighbors around the TMDs in different configurations (monomeric, or dimeric in righthanded ***χ***(−) and left-handed ***χ***(+) configuration) in the symmetric (left) and asymmetric (right) bilayer compositions and the respective ratios between the average and the variance of the residence times (bottom).

Overall, the residence times of PC and ERG do not differ considerably. They are slightly increased for dimers and do not show differences between the symmetric and asymmetric bilayers. Surprisingly, the residence times for ERG do not differ between the extracellular and the cytosolic side either. Only the residence times of PS lipids are roughly twice as large as for PC lipids, which is congruent with the notion of strong electrostatic interactions between the charged PS head groups and the R/H residues of the Slg1 TMD. ERG shows an increased average residence time in the asymmetric bilayer around the ***χ*** (−) dimer and around the monomers, despite not being enriched in these configurations. The ratios of the mean and standard deviations of ***τ*** lie between 2 and 4, and, accordingly, the lipid residence times do not follow a simple exponential decay. This observation applies to all lipid species and appears to be a general property of the TMD-lipid contacts.

Therefore, the ERG-TMD binding cannot be considered a strong interaction. Unlike the PS-TMD bond, the ERG-TMD bond is transient, and the reported enrichment of ERG implies a high rate of molecule exchange around the TMDs.

### 3.2 Characterization of the TMD geometry within the bilayers

In this section, we elucidate the position of the TMDs within the membrane characterized by the TMD tilt and the distance of selected residues to the bilayer center.

We show the tilt angle distribution of each TMD in figure 5, with the tilt defined as the angle between the vector of respective COMs of the second and second to last three C_α_ carbons to the bilayer normal (z-axis).

**Fig. 5.**
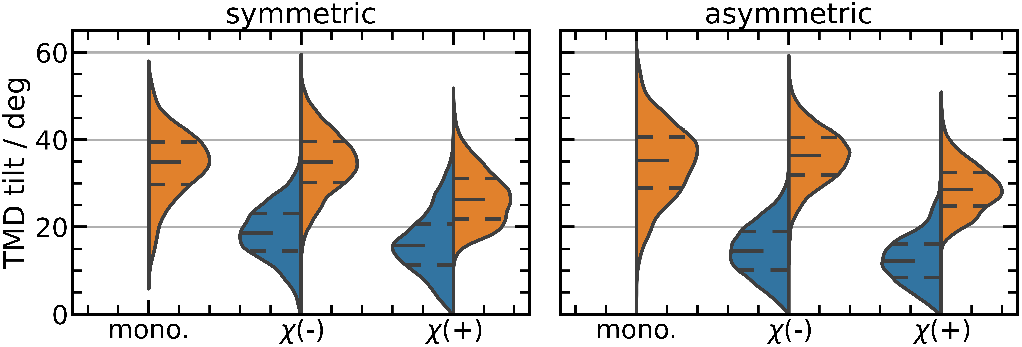
The TMD tilt angle in the symmetric (left) and asymmetric (right) bilayer composition for each TMD configuration (monomer, dimer ***χ***(+) and ***χ***(−)). Orange and blue indicate the tilt values of the TMD with higher and lower tilt, respectively.

The tilt distributions display a similar pattern for all configurations and compositions, with one TMD exhibiting a high tilt angle of roughly 35° (and 25° for the ***χ*** (+) dimer) and the other TMD showing a lower average tilt of approximately 15° to 20°. In some details, the tilt distributions differ: First, the ***χ*** (+) dimers exhibit an overall lower tilt angle and a lower discrepancy between the TMDs. Second, the tilt distributions of TMD monomers are somewhat broader than that of the dimers, and third, the average tilt is slightly decreased in the asymmetric bilayers.

We calculated the correlation between the tilt angles of the TMDs in the dimers listed in table 1 to characterize the rigidity of the dimer bonds. Limiting values of the correlation coefficients can be showcased by a rigid X-shaped dimer of two sticks. Corresponding correlation coefficients of the tilt angle would be −1 for a tilt within the X-plane and 1 for tilting the X-plane itself, while the flexibility of the bond reduces the absolute correlation.

**Table 1.**
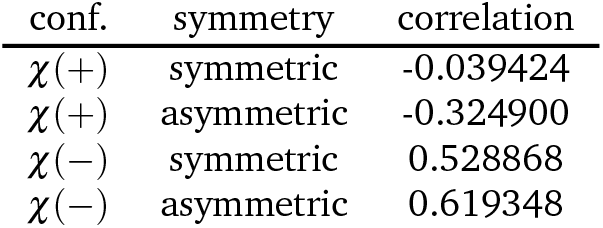
Pearson correlation coefficients between the tilt angle of the two TMDs in each dimer.

The TMDs in the ***χ*** (−) dimers exhibit a positive correlation, whereas the correlation coefficient is small and negative for the ***χ*** (+) dimer in the asymmetric bilayer. In the symmetric bilayer, we find no correlation between the ***χ***(+) TMDs. This is a strong indication that the ***χ*** (−) dimers display a signifcantly stronger interaction which is particularly for the asymmetric (and thus more realistic) bilayer. Further below, this conclusion will be corroborated by the evaluation of the crossing angle distributions.

With the tilt angle, the z-position of the motifs within the bilayer may change. We, therefore, show the distribution of distances to the bilayer centers of residue G9 in comparison with the distance distribution of nearby *(r_xy_* ≤ 0.6 nm) ergosterol molecules in figure 6). We chose G9 as a representative for a high affinity GxxxG contact (compare figures S1^†^ and S2^†^). For the dimers, the height distributions of the two respective G9 residues are averaged.

**Fig. 6.**
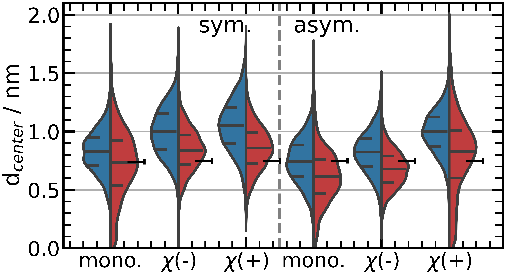
Distributions (violin representation) of the average distance of residue G9 (blue) and ERG TMD-neighbors (within 6 Å of a backbone C_α_) (red) to the bilayer center in the extracellular leaflet. Lines within the violins indicate the quartiles of the respective distributions. The black horizontal lines depict the position of the 50% quartile of bulk ergosterol distances, while the attached vertical lines indicate the positions of the 25 and 75% quartiles (representing the distribution width).

As expected, the variation of the average distance of G9 is correlated with the tilt angle (compare figures 5, 6). The ***χ*** (+) case, displaying the lowest tilt angle, displays the largest distances. Closer inspection reveals that for the three cases (monomer,***χ*** (−), ***χ*** (+)) the distances for the asymmetric case are lower by nearly 0.1 nm, reflecting an overall shift of that residue due to the lipid composition. For all cases, the ERG center of mass lies slightly beneath G9 (approx. 0.1 nm). Remarkably, this shift is basically independent of the affinity as expressed by the enrichment factor and independent of the average position of all ERG molecules in the respective systems. In contrast, the width of the distribution indeed reflects the TMD-ERG affinity. While, the ERG distribution is significantly increased next to the TMDs compared to the bulk ERG molecules, in both cases without any enrichment, namely the monomer in the symmetric bilayer and the asymmetric ***χ*** (+) display a much broader distribution for ERG. This points towards a specific impact of the strength of the TMD-ERG interaction on the ERG position.

Similarly, we show the distance distribution of residues R29/H30 in comparison with the distance distributions of the phosphor atoms of nearby phospholipids and, again, the distance distributions of the R/H residues in the dimers are averaged in figure 7. From this representation, we can evaluate the effect of charged PS-R/H interactions on the distances of the TMDs and bilayer thicknesses.

**Fig. 7.**
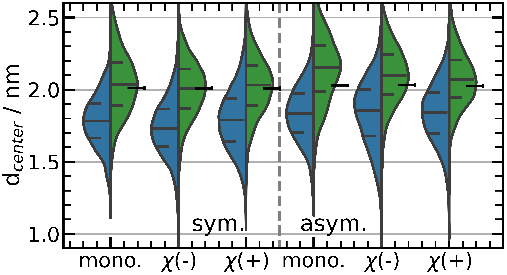
Distributions (violin representation) of the average distance of residues R29/H30 (blue) and phosphor atoms of nearby phospholipids (within 6 Å of a backbone C_α_) (green) to the bilayer center in the cytosolic leaflet. Lines within the violins indicate the quartiles of the respective distributions.

The heights of the residues are basically independent of the TMD state. Similar to the observations in the extracellular leaflet, though, we see a small but systematic shift when comparing the symmetric and the asymmetric case. Furthermore, the distances of the phosphor atoms of the phospholipids are sensitive to the lipid composition. Whereas in the symmetric case they are hardly influenced by a nearby TMD, in the asymmetric case the phospholipids close to the TMD are significantly shifted away from the membrane center as compared to an average phospholipid. Furthermore, this effect is most prominent for the case of the strongest affinity (monomer, asymmetric case). Thus, as a second difference to the behavior at the extracellular side, a stronger affinity between TMD and lipids does not narrow down the distribution of distances but rather shifts the total distribution. Naturally, this shift gives rise to local variations of the thickness of the membrane.

### 3.3 Characterization of the TMD contact

The distribution of the crossing angles of the TMD dimers can give information about the flexibility of the TMD dimer bond. We depict the crossing angle distribution for each dimer simulation in figure 8. The crossing angle was calculated according to ref.^68^, which is defined as the dihedral angle of the beginning of the helices (COMs of the fist three C_α_ carbon atoms of each helix) and the two points, which are at the closest distance on the two vectors assigned to the helices.

**Fig. 8.**
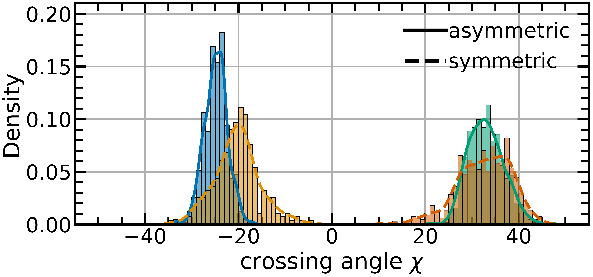
Crossing angle distributions of the TMD dimers in right-handed ***χ***(−) and left-handed ***χ***(+) configuration.

The crossing angle distributions are narrower in particular for the ***χ*** (−) configuration. Furthermore, there is a slight but significant effect of the composition (more narrow in the asymmetric case). This fully agrees with our interpretation of the tilt correlation coefficients, namely higher bond stability for a narrower crossing angle distribution.

To further characterize the bond of the dimer, we recorded snapshots of structures of the TMD dimers, shown in figure 9. From visual inspection of the snapshots, one can obtain information on the nature of the TMD contacts. In the ***χ***(+) configuration, the motif faces of the two TMDs are oriented anti-parallel (black spheres on opposite sides), while the TMDs are parallel in the ***χ***(−) configuration. Müller and coworkers^69^ could show that the optimal orientation of the glycophorin dimers depends on the sign of the crossing angle and, thus, the different crossing angle configurations naturally imply different orientations. The dimer configurations that show a high affinity to ERG (***χ*** (−) and ***χ*** (+), symmetric) exhibit one similarity: The TMD contacts are formed via motif a) and b) (green and red) or, in the case of ***χ*** (−) in the symmetric bilayer motif b), with another contact face (green and gray), such that the respective “free” motif faces form a broad contact surface on one dimer side. For ***χ***(+) in the asymmetric bilayer that shows no ERG affinity, motifs a) (red) are in contact such that the free motif faces (green) lie on opposite dimer sides. This configuration likely behaves similar to the monomers, and indeed, the monomers also do not show extensive affinity to ERG molecules.

**Fig. 9.**
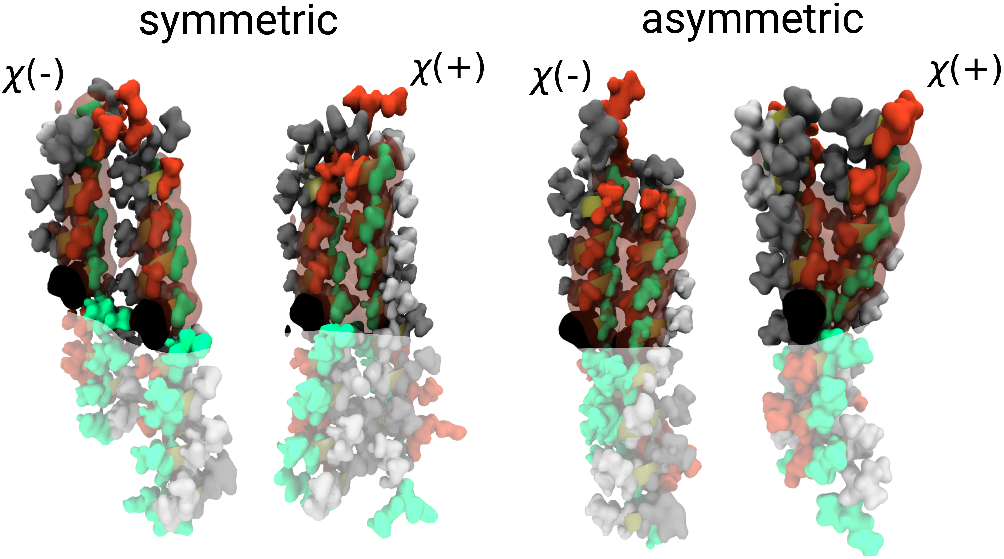
Snapshots of representative configurations for each dimer configuration and bilayer composition. Residues highlighted in red and green form the TMD faces including motifs a and b, respectively. The black spheres highlight the positions of residue 16 in the center of the sequence, and the red shading indicates the position of glycine residues. The shaded area depicts the cytosolic side of the TMDs.

Furthermore, we quantified the contact closeness represented by the sums of contacts between respective TMD faces per simulation and show the results in in table 2. The sums indicate a significant impact of the lipid composition on the closeness of the TMD contact, as they are systematically increased in the asymmetric bilayers. Especially, the sum of contacts per simulation is high for the ***χ*** (−) dimer in the asymmetric bilayer.

**Table 2.** Sum of the contact fractions shown in figure S4^†^ for each composition and TMD configuration.

| symmetry | conf. | contact sum (extrac.) | (cytos.) |
| --- | --- | --- | --- |
| asymmetric | $\chi(-)$ | 6.724209 | 0.487437 |
| asymmetric | $\chi(+)$ | 2.390279 | 0.421235 |
| symmetric | $\chi(-)$ | 1.581914 | 0.286553 |
| symmetric | $\chi(+)$ | 1.468211 | 0.761027 |

Therefore, the ***χ*** (−) and ***χ*** (+) configurations exhibit distinctive contact profiles. The profiles conform to the notion that the lefthanded contact via the GxxxG motif is energetically favorable^69^ and may indicate that the bilayer asymmetry (presence of charged lipids) favors a tighter packing of the dimers. The contact profiles of the ***χ*** (+) dimer in the asymmetric bilayer might explain the lack of ERG enrichment: The contact surface is somehow blocked by the protein contact.

### 3.4 Comparison to coarse-grained simulation data

In this section, we compare a selection of our results from allatom (AA) simulations to the corresponding CG simulations. This comparison can help to assess the similarity but also differences between the two representations.

In an initial step we intend to check whether the AA structures, obtained from the CG structures via the mapping procedure, are close to equilibrium. For this purpose, we computed the RMSD between the initial CG structure and remapped (AA to CG) structures from our atomistic simulations. The time series of the RMSD values is depicted in figure 10. The initial RMSD at time 0 is the RMSD that is inherent in the backmapping process and lies roughly between 1.5 and 2.5 Å. Already after 500 ns the RMSD reaches a plateau for all systems. Interestingly, the average RMSD of two dimer structures, recorded in pure CG simulations after time intervals of the order of microseconds, display very similar values. This is a strong indication that dimer structures after mapping from CG to AA are indeed close to the equilibrium AA structures. Otherwise, extensive reorganization of the AA structures would have brought the system in different regions of the phase space, which would still be visible after final remapping to the CG level and comparison with the initial CG structure. In one case, though, the RMSD values show slightly higher differences. The RMSD of the atomistic ***χ*** (−) symmetric bilayer is highest, while it is the lowest in the CG representation. However, these differences are small (≈ 1 Å).

**Fig. 10.**
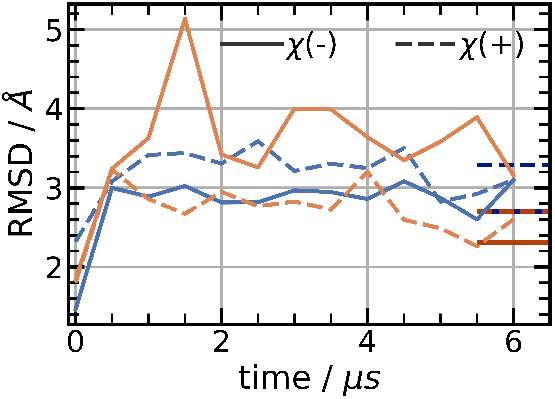
Time evolution of the RMSD in atomistic simulations. Blue and red indicates data from the asymmetric and symmetric bilayers, respectively. The horizontal lines in darker color depict the average RMSD values for random structures pairs in the CG simulations.

Since the CG structures are indeed reasonable, we may now start with a closer comparison between the AA and the CG configurations. In figure 11 we present the lipid enrichment near the TMD faces, the lipid neighbor residence times, and the crossing angle distributions for the CG simulations. Starting with the comparison of the enrichment of ERG near the extracellular faces of the TMDs (figure 11, top), we see that also in the CG simulations ERG is enriched near the motifs and significantly less enriched at the other sites. However, also deviations between the AA and the CG simulations are present. First, the separation between the motif and the other sites is weaker, second, the non-binding character of ***χ*** (+) in the asymmetric case is not reproduced. For PC, the observed absence of enrichment of PC is similar between AA and CG. Looking at the enrichment near the cytosolic TMD faces (figure 11, middle left), the ERG affinities, obtained via CG, are similar to the reported affinities in the extracellular leaflet. Again, this contrasts with the atomistic results where ERG shows no enrichment. For PS, just like for PC lipids, the enrichment data matches. The corresponding data for ERG and PC enrichment in the symmetric bilayer show similar behavior (compare figure S5^†^, right). In the Martini force-field, diffusion constants are increased compared to diffusion constants of atomistic simulations. Therefore, we expect the residence times to be shorter in the CG representation, and, indeed, for PC, the residence times are reduced by roughly two thirds with a simple exponential decay as indicated by s(***τ***)/ < ***τ*** >≈ 1 (figure 11, bottom). For PS, this decrease is slightly stronger with a factor of 2. In contrast, for ERG, the residence times are similar to those from the atomistic simulations, which reflects the overall elevated ERG affinity. Furthermore, the crossing angle distributions do not show a significant composition dependence and exhibit two maxima for composition, whereas the distributions show distinct maxima in the atomistic simulations. This difference presumably is related to the RMSD difference of the ***χ*** (−) symmetric bilayer. When comparing the contact sums that quantify the closeness of the dimers, the lack of any PS effects is also visible. Here, the contact sums in the CG are all very similar, whereas in the atomistic simulations the contact sums were systematically higher in the asymmetric bilayers (compare tables 3) and 2)).

**Fig. 11.**
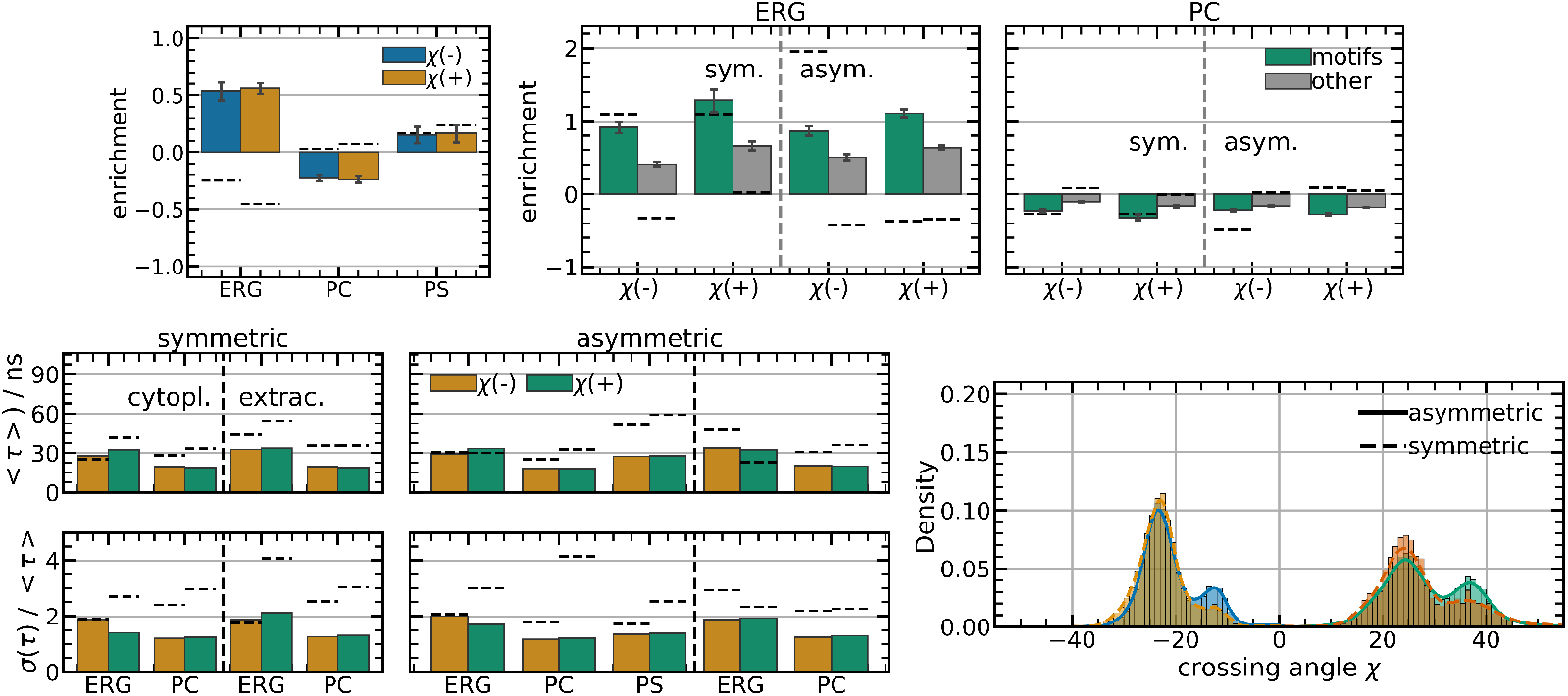
Evaluated data from the atomistic simulations (dashed horizontal lines) compared to the corresponding data from CG (Martini) simulations, namely, enrichment of lipids in the cytosolic (top left) and the extracellular leaflet (top right), lipid neighbor residence times (bottom left), and the crossing angle distributions (bottom right).

**Table 3.** Sum of the contact fractions of the TMD dimers in the CG simulations shown in figure S5^†^ for each composition and TMD configuration.

| symmetry | conf. | contact sum (extrac.) | (cytos.) |
| --- | --- | --- | --- |
| asymmetric | $\chi(-)$ | 3.929107 | 2.251475 |
| asymmetric | $\chi(+)$ | 2.241676 | 2.183482 |
| symmetric | $\chi(-)$ | 3.947805 | 2.248775 |
| symmetric | $\chi(+)$ | 3.741326 | 1.901210 |

Thus, the properties of the system are congruent on a qualitative level but differ in the details. The enrichment data and expected residence times for the phospholipids of the CG match the atomistic data very well. The ERG enrichment data is systematically overestimated in the CG simulations, indicating an exaggerated TMD-ERG interaction. As the enrichment data only deviates for ERG, we suggest that this deviation is not due to the limited sampling in AA representation but an artifact of the Martini force-field. The interpretation of the deviations between the TMD bond is more complicated. As we have shown that the dimer bond is very strong and inflexible, especially in the right-handed configuration, insufficient sampling of the dimer structure space make the comparison between simulations harder, even though we suspect to have sufficiently sampled the configuration space within the > 6.5 μs simulations. This assumption is corroborated by the time series of the RMSD values that are more or less converged to a plateau value. The crossing angle distributions as well as the contact sums indicate that the behavior of the CG TMD dimers indeed differs from the behavior of the dimers in atomistic simulations. We particularlz stress the insensitivity to the lipid composition on the TMD bond in CG resolution (lack of PS effect). Therefore, we attribute these differences to the lack of molecular details in the Martini representation. It has been shown that the protein-protein interactions are overestimated in Martini 2.2^70,71^, as also explicitly seen in our previous work^38^. Therefore, it might be possible that also the protein-lipid interactions suffer from a similar overestimation. However, since currently there is no sterol force field for Martini 3, reproduction of the current results with Martini 3 is not possible.

## 4 Conclusion

In this work, we studied the impact of the lipid environment on different TMD configurations of the Slg1 receptor via atomistic simulations and compared it with corresponding simulations on the coarse-grained level.

The presence of charged PS lipids changed the bilayer thickness and the height of ERG molecules near the TMDs. Ergosterol exhibited a high affinity to the GxxxG motifs of the TMD dimers in the extracellular leaflet. On the other hand, we could not observe a substantial enrichment of ergosterol near the TMD monomers and, consequently, the presence of two TMDs with their motifs aligned was necessary for an increased affinity to ERG in this model system. The lack of ERG enrichment at the GxxxG motifs of the ***χ*** (+) dimer could be related to its distinct TMD contact, where the motifs faced towards each other and, in effect, were partially screened from lipid contacts.

The comparison between atomistic and coarse-grained simulations was particularly helpful for the characterization of the TMD contacts. Due to the TMD bond strength in CG simulations, sampling the relevant configurational space of the dimers is particularly challenging. Therefore, we validated the sampling of the dimer-containing systems in atomistic representation and showed that the RMSD of the dimer complex reaches a plateau already within 500 ns that is stable for the residual 6 μs. Even though the protein-protein interaction is reportedly too strong in the Martini force field^70,72,73^, the crossing angle distributions of the CG simulations showed multiple maxima (as compared to one maximum in AA) and especially the crossing angle distributions of the ***χ*** (+) dimers differed between the AA and CG description. These differences are also reflected comparing the closeness of the TMDs. In the AA simulations, the closeness of the TMD contacts showed high sensitivity on the lipid composition: The sum of contacts was systematically increased for the dimers in the asymmetric bilayers, especially for the ***χ*** (−) dimer. In the CG simulations, we did not observe any impact of the lipid composition on the TMD dimer bond. We attributed these differences to the mentioned shortcomings of the protein-protein interaction description in Martini 2. With regards to the TMD-lipid affinities measured by the lipid enrichment factor, we conclude that the Martini forcefield could very well reproduce the properties of the PS and PC lipids, while the TMD-ERG affinities were systematically overestimated. Martini 3.0 has significantly improved the description of proteins and reportedly mended artifactual interactions. ^40^ Future studies will have to develop new force fields for sterols and validate their behavior in different membrane environments.

In our previous work^38^ we had concluded from the CG simulations, that the stability of the dimer formation upon addition of PS hardly changes. This was extracted from the insensitivity of the Gibbs free energy of dimer formation as well as the crossing angle distribution when adding PS, i.e. when comparing the asymmetric with the symmetric case. This agreed well with the fact that the experimentally obtained network factor hardly changed as well. In contrast, the present data suggest that ***χ*** (−) is more stable in the asymmetric case, based on the large value of the contact sum and its quite narrow distribution of crossing angles. This prompts the question whether for the understanding of the network factor processes beyond the dimer formation matter. Note that the network factor can only be measured with a resolution of >100 nm such that cooperative effects and complex formations of multimers would contribute. This work can act as the foundation to further study the characteristics of the formation of trimers or multimers using a similar multiscale approach. The utilization of the Martini 3 force field in the longer run may also lead to an increased fidelity of description and improve the calculations of the free energies of multimer formation that are otherwise inaccessible in atomistic resolution.

In summary, we characterized the interaction sites of the Slg1 TMD by a systematic study of TMD-lipid and TMD-TMD interactions, using a multiscale approach. The findings may act as the basis for future studies regarding the effect of the juxtamembrane regions or even ectodomains and may shed light on the signaling mechanism of the Slg1 stress sensor.

## Conflicts of interest

There are no conflicts to declare.

## Acknowledgment

We acknowledge the Deutsche Forschungsgemeinschaft (DFG) for funding via SFB 1348. Furthermore, we appreciate very helpful correspondence with S.J. Marrink and P.C. Souza.

## Notes

### Competing Interest Statement

The authors have declared no competing interest.

