## supplemental_information for "Impact of bilayer composition on the dimerization properties of the Slg1 stress sensor TMD from a multiscale analysis ^†^"

### ESI

The following files are available free of charge.

**Table S1** System, simulation lengths and the compositions of the CG simulations.

| System | len. ($\mu$s) | #DOPC | #POPC | #DOPS | #POPS | #ERG | |
|---|---|---|---|---|---|---|---|
| $\chi(-)$, symmetric | 10 | 224 | 112 | 0 | 0 | 0 | 84 |
| $\chi(-)$, asymmetric | 10 | 192 | 96 | 32 | 16 | 84 | |
| $\chi(+)$, symmetric | 10 | 224 | 112 | 0 | 0 | 0 | 84 |
| $\chi(+)$, asymmetric | 10 | 192 | 96 | 32 | 16 | 84 | |

**Table S2** Simulation lengths, temperatures and compositions of the utilized MD simulations.

| System | len. ($\mu$s) | #DYPC | #POPC | #YOPC | #DYPS | #POPS | #YOPS | #ERG |
|---|---|---|---|---|---|---|---|---|
| $\chi(-)$, symmetric | 6.81 | 112 | 112 | 112 | 0 | 0 | 0 | 84 |
| $\chi(-)$, asymmetric | 6.76 | 96 | 96 | 96 | 16 | 16 | 16 | 84 |
| $\chi(+)$, symmetric | 6.77 | 112 | 112 | 112 | 0 | 0 | 0 | 84 |
| $\chi(+)$, asymmetric | 6.91 | 96 | 96 | 96 | 16 | 16 | 16 | 84 |
| monomer, symmetric | 1.33 | 138 | 139 | 138 | 0 | 0 | 0 | 106 |
| monomer, asymmetric | 2.0 | 121 | 122 | 121 | 18 | 18 | 18 | 106 |

**Table S3** Ensembles and force constants used during the equilibration steps on head group positions and chain dihedrals.

| step | time / ps | ensemble | P atom z pos. (kJ/mol) | chain dih. ang. (kJ/mol) |
|---|---|---|---|---|
| 1 | 25 | NVT | 1000 | 1000 |
| 2 | 25 | NVT | 1000 | 400 |
| 3 | 25 | NVT | 400 | 200 |
| 4 | 100 | NpT | 200 | 200 |
| 5 | 100 | NpT | 40 | 100 |
| 6 | 100 | NpT | 0 | 0 |

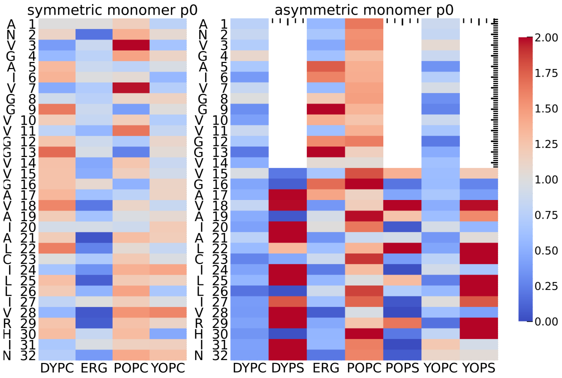


**Fig. S1** Enrichment and depletion maps of lipid neighbors around the TMD monomer residues in the symmetric (left) and asymmetric (right) bilayer. The number of lipid type neighbors is normalized with respect to the total number of neighbors and the respective fraction of the lipid type within the leaflet, such that values above unity indicate enrichment and values below unity indicate depletion of the lipid type. Red indicates values of 2 or higher.

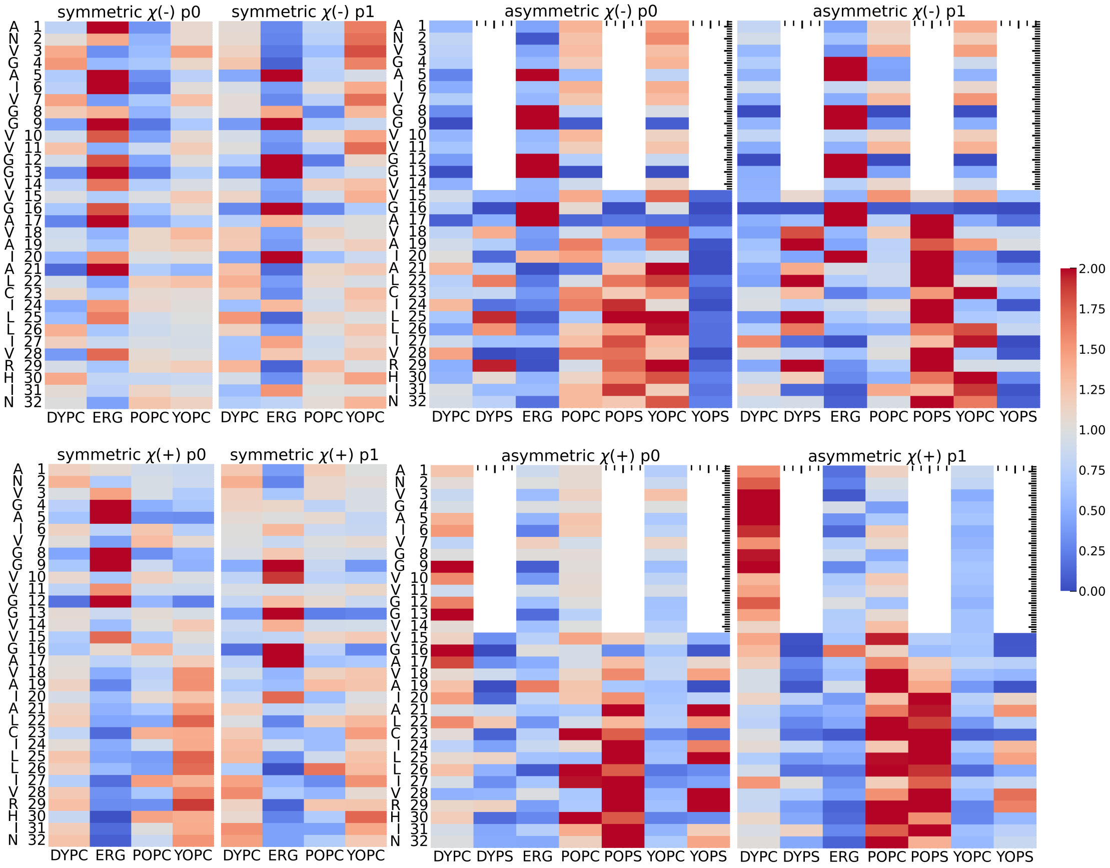


**Fig. S2** Enrichment and depletion maps of lipid neighbors around each TMD (p0 and p1) in the dimer configurations with negative crossing angle, $\chi(-)$ (top), and positive crossing angle, $\chi(+)$ (bottom), in the symmetric (left) and asymmetric (right) bilayers. The number of lipid type neighbors is normalized with respect to the total number of neighbors and the respective fraction of the lipid type within the leaflet, such that values above unity indicate enrichment and values below unity indicate depletion of the lipid type. Red indicates values of 2 or higher.

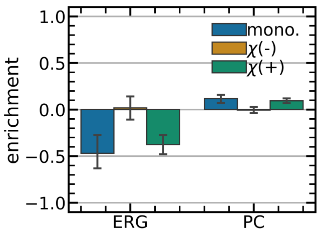


**Fig. S3** Enrichment of ERG and PC lipids in the cytosolic leaflet of the different symmetric bilayer setups (monomeric, or dimeric in right-handed $\chi(-)$ and left-handed $\chi(+)$ configuration) around the TMD. The enrichment is defined as the number of the respective lipid species divided by the total number of neighbors and the total concentration of the species minus 1 such that 0 indicates no enrichment or depletion.

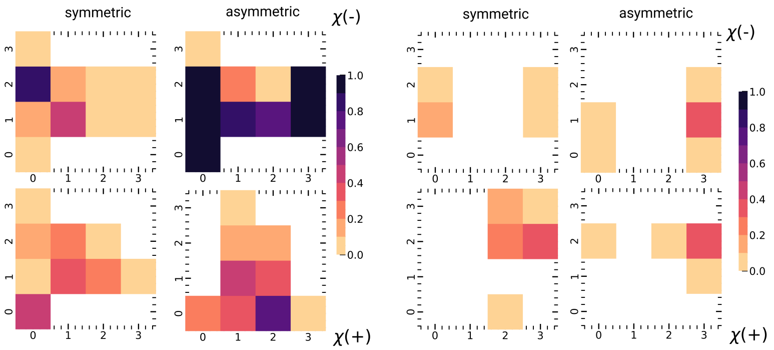


**Fig. S4** The TMD-TMD contact map of the extracellular (left) and cytosolic (right) TMD faces. Indexes 0 and 1 indicate the TMD face including motifs a) and b) and indexes 2 and 3 indicate the other TMD faces. The color code represents the fraction of contacts (distances of less than 7Å) formed per simulation such that a value of 1 represents a contact that was maintained throughout the whole simulation. The distance cutoff was chosen such that at least one contact pair in all bilayer systems exhibits a contact fraction of 0.9.

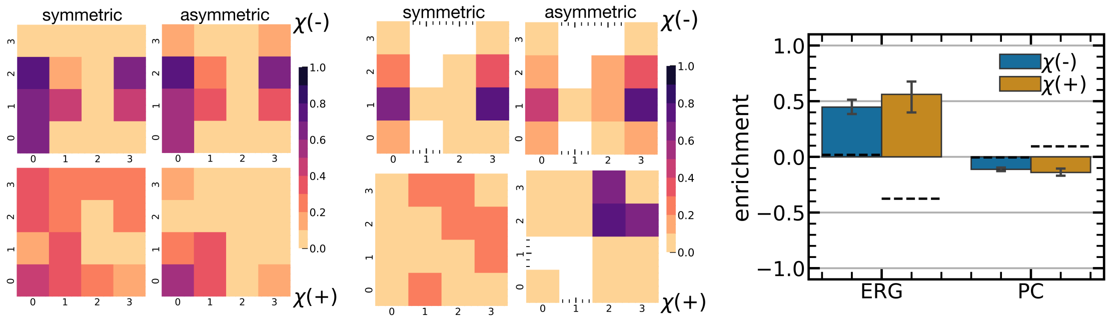


**Fig. S5** The evaluated data from the atomistic simulations (dashed lines) compared to the corresponding data from coarse-grained (Martini) simulations, namely, enrichment of lipids in the cytosolic leaflet of the symmetric bilayer (left), and the protein contact map of the cytosolic TMD side (right).
